# Information content of downwelling skylight for non-imaging visual systems

**DOI:** 10.1101/408989

**Authors:** Ryan Thiermann, Alison Sweeney, Arvind Murugan

## Abstract

Light-sensitive proteins (opsins) are expressed in non-imaging tissues like the brain, dermis and reproductive organs of most animals. Such tissues have been shown to sense the intensity and spectrum of light over time. Functional links to circadian and reproductive rhythms have been speculated but remain uncertain. Here we use information theory to quantify the ‘natural scene’ for non-imaging opsins, i.e., spectral patterns in downwelling skylight. Our approach synthesizes measurements of natural downwelling spectra, atmospheric distortions, and weather, with the biophysical constraints of opsins and biochemical clocks, while minimizing assumptions about how organisms process such information. We find that tissues expressing multiple opsins could use twilight to extract significant information about lunar phase and time of day in many climates. In contrast, information in light intensity is far less robust to atmospheric perturbations. Thus our work quantifies circalunar and circadian regularities in the spectrum of downwelling radiance salient to non-imaging opsins.

## I. INTRODUCTION

Vision using imaging eyes is a remarkable product of evolution[1, 2]. However, numerous non-ocular tissues are also known to express photosensitive proteins – opsins that are close evolutionary variants of the visual opsins used in the retina[3–6]. The planar geometry of these diverse tissues precludes imaging vision, but such tissues can still detect the intensity of light and potentially resolve its spectrum. In fact, it has been argued [7] that such non-imaging tissues can be very sensitive to low intensity light, e.g., down to the intensity of starlight. In contrast, the focusing geometry needed for imaging vision must necessarily discard significantly more light and thus be far less sensitive.

Extraocular opsins are found extensively in vertebrates often in brain and reproductive tissues - and in numerous marine organisms like corals, squids, barnacles and starfish [7–10]. The biological role for these opsins is not clear and is being actively investigated[8, 11–14]. While the expression of some of these opsins could be an evolutionary “spandrel” and not currently serve any function, disruption of natural inputs to some of these opsins are being increasingly implicated in a variety of physiological conditions and diseases[15–22], ranging from disruption of testicular and other sexual development in birds[23, 24], circadian[25] and seasonal rhythms in reproduction [13, 26, 27] to metabolic and other such disorders [5, 15, 28].

Hence it is important to understand the ‘natural scene’ — i.e., the statistical structure of natural light stimuli — that has been present over evolutionary timescales for these non-visual imaging systems. Regularities and patterns in natural scenes can point to possible functions and functional adaptations. For example, the natural scene for imaging vision has been characterized, e.g., in terms of edges and corresponding edge detectors in the retina[29–33] while recent work has focused on how natural scenes have shaped other senses such as the auditory system and olfaction[34–36].

The natural ‘scene’ for non-imaging vision can be expected to be distinct from imaging vision in two important ways - (1) while lacking the spatial structure associated with images, the scene can involve rich temporal patterns associated with the natural world, (2) the spectrum of light can be expected to play a critical role since these can be measured to great accuracy by comparing the photon catch in opsins with different spectral sensitivity [17, 37, 38].

Here, we characterize the regularities and irregularities in the natural stimuli available to non-imaging visual systems. We use an information theoretic approach that quantifies the biophysically resolvable patterns without assuming specific downstream internal processing in an animal’s neural or other clock-like systems. This approach provides an upper limit on the robustness of physical patterns available to non-imaging visual systems. Similar approaches have recently been used to understand temporally coded information in gene regulation[39].

We find two strong regularities in downwelling natural light: a circadian cycle in downwelling spectrum with a sharp feature at twilight, and a circalunar modulation of this circadian rhythm. Such regularities in spectrum are robust when spectral distortions due to weather are taken into account and perceptible given biophysical constraints inherent to opsins. However, the correlated regularities in light intensity are not robust to these factors because they are overwhelmed by weather-related fluctuations.

While numerous natural signals are available and have been quantified for biological rhythms synchronized to the period of a day (circadian) or the period of a year (circa-annual), few widely available and reliable signals are known that could synchronize such 28-day circalunar rhythms. However, important biological phenomena, notably reproductive phenomena, have synchronized components that depend on this period. Many organisms, such as echinoderms, cnidarians and corals show reproductive behaviors at specific points of the lunar cycle [16, 17, 26, 27, 41], including the dramatic synchronized mass spawning in corals and other marine invertebrates; humans also show 28-day periodicity of the reproductive cycle.

Our results therefore quantify a novel information-bearing signal - i.e., a *zeitgeber* - potentially relevant to physiology with circalunar periodic features. Our work suggests that non-imaging opsin-based visual systems are well-suited to perceiving the robust circalunar rhythms coded in the spectrum of natural light.

## II. RESULTS

### A. Non-imaging opsins can sense temporal patterns in spectrum at naturally-occurring intensities

The geometry of non-imaging tissues allows sensing of light to much lower intensities than the focusing geometries needed for imaging vision [7]. See Fig.1a,b. As a result, even the small amounts of light that penetrates e.g., the skull is likely enough to be sensed by tissues such as deep brain tissue in birds or a developing mouse[8]. The likelihood of light at these naturally occurring intensities is detectable within tissues increases given that initial evidence indicates that non-imaging opsins also tend to have very long (minutes or hours) integration times [3].

**FIG 1.**
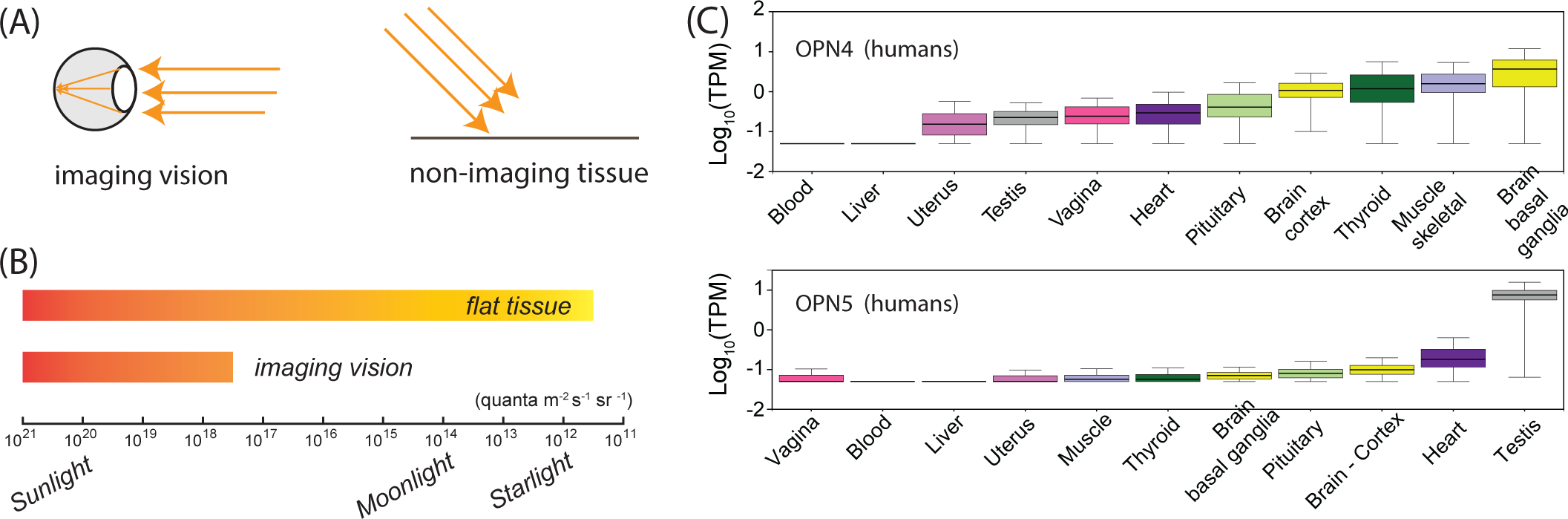
Light-sensitive opsins, expressed in numerous non-imaging tissues, can sense light of much lower intensity than imaging eyes. (A,B) Flat tissue with light sensitive proteins (opsins) can sense light down to intensities comparable to that of starlight. In contrast, imaging vision requires focusing geometries that reduce received intensity and are thus less sensitive. (Data in (B) reproduced from [7]). (C) Non-imaging opsins are expressed at significant levels in non-occular tissues in humans (data from GTEx Portal[40]) and numerous land and marine animals; disruption of light input to these tissues can disrupt circadian, reproductive or other behaviors. In contrast, rhodopsin, used for imaging vision, is mostly expressed in the retina (not shown). (Transcripts-Per-Million (TPM) quantifies the fraction of total transcripts present[40] in that tissue.)

In humans, the non-visual opsins melanopsin (OPN4) and neuropsin (OPN5) are found to be expressed in nu-merous non-occular tissue. An example is shown in Fig.1c. Similar results have been reported in other organisms[8]. In contrast, the same dataset shows that rhodopsin is primarily expressed in the retina.

Any functions of extraocular opsin expression are not yet clear. Obviously, colonies of lab animals remain viable over many generations under artificial illumination lacking any of the spectral features we describe here. At the same time, numerous anomalies have been reported when natural illuminants are disrupted. These disruptions range from sexual development in ducks being inhibited by placing a black cap on the head [42] and disruptions in seasonal reproduction in various other birds [8, 11] to hyperactivity in zebrafish larvae [9, 12]. Similarly, the advanced material culture of humans may buffer historical or pre-historical disadvantages to suboptimal timing of reproduction [43].

Because there is no structure for imaging in non-ocular tissues, the ‘natural scene’ for such non-imaging opsins cannot involve spatial structure. The intensity and spectrum of the light could be expected to play a role, since these non-visual opsins typically have distinct spectral sensitivities [10]. Further, the intensity and spectrum can be observed over time. Thus, we define the ‘natural scene’ for non-imaging opsins as changes in the spectrum and intensity of natural light over time.

### B. Daily and monthly rhythms in the spectrum of natural light

The most obvious-to-humans systematic variation in natural light is the change in intensity over 24 hours, due to the rising and setting of the sun. However, as the sun changes elevation, the path length of direct solar light through the atmosphere changes; since the atmosphere absorbs more in some wavelengths than others, and the magnitude of wavelength-dependent scattering increases, the downwelling spectrum changes. These relative changes are minor when the sun is above 20 degrees in elevation, but dramatic at evening twilight and at dawn. Fig.2b shows the systematic variation in the relative amount of light at 440 nm and 550 nm in the hours after sunset on a new moon night (which we label Day 14); the curve shown is a fit to experimental observations[44] from rural Pennsylvania, USA.

**FIG 2.**
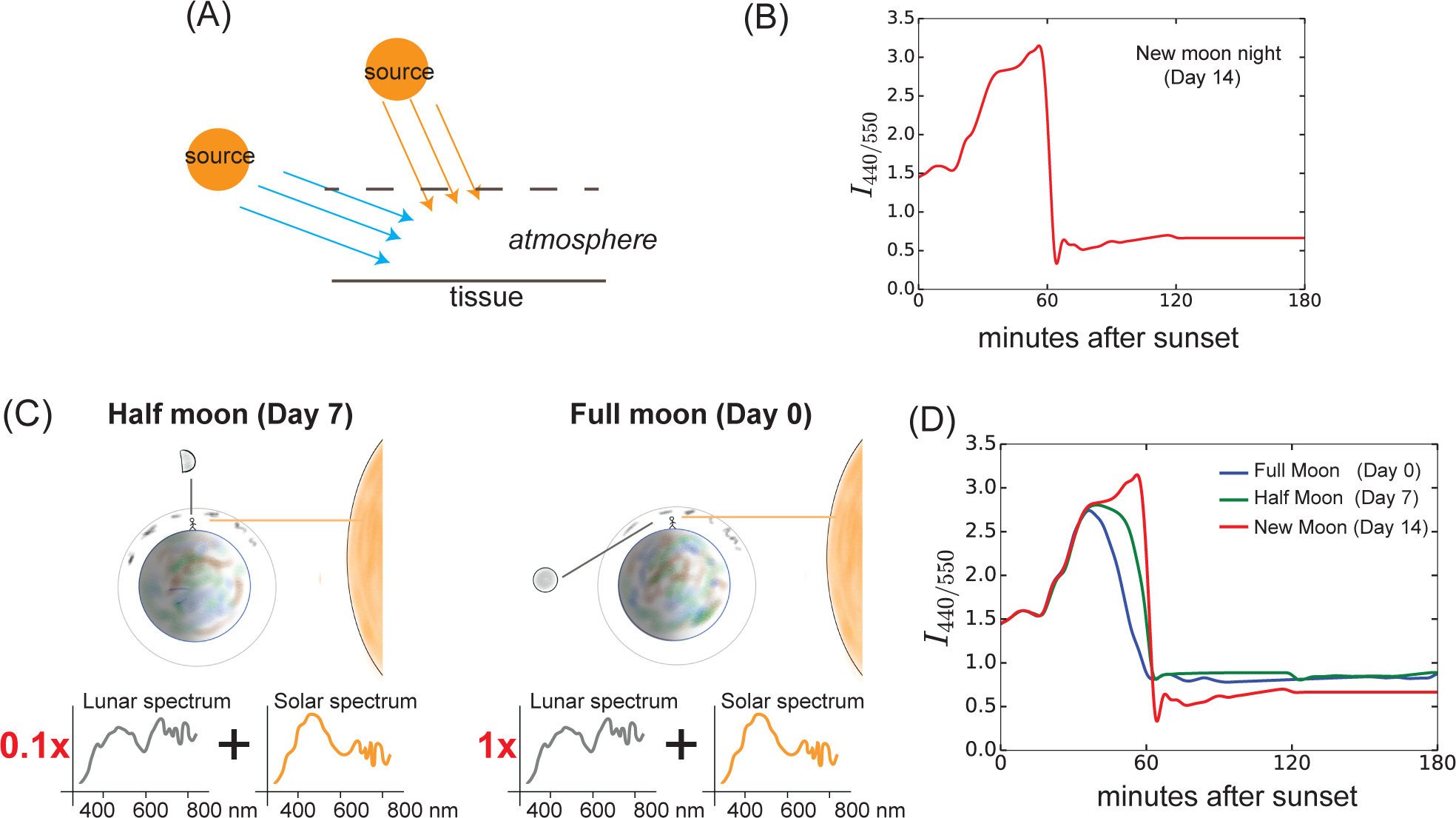
The strongest regularities in the spectrum of downwelling natural light are a daily rhythm due to the setting (or rising) of the sun and a monthly modulation of this daily rhythm due to the rising (or setting) of the moon. (A) The changing elevation of the sun causes dramatic spectral changes at twilight. (B) Hence the ratio of light as captured by two different opsins with peak sensitivities at, say, 440 and 550 nm, shows a specific dynamic pattern during twilight hours. (c) The moon, whose spectrum is distinct from the sun, contributes and thus changes the spectrum of twilight (and dawnlight). However, the strength of the lunar contribution is modulated over the 28-day lunar cycle. (d) Hence the time course of twilight spectrum potentially carries information about the lunar phase. (Panels (b,d) based on measurements in [44].)

Consequently, the downwelling spectrum over twilight hours changes with the specific night of the lunar cycle. See Fig.2c,d (data from [44], also observed in [17, 37]).

However, atmospheric variations, such as cloud cover and humidity, can significantly change the actual intensity and spectrum of light reported in Fig.2 (see Supplementary Fig. 1).

The most significant distortion in these patterns of downwelling natural light is due to variable cloud cover. For example, cloud cover can dramatically reduce the transmitted intensity; as shown in Methods based on data in [45], transmitted intensity can be reduced by more than an order of magnitude as cloud cover varies from clear skies to completely overcast. In contrast, the impact of cloud cover on spectrum is much smaller since clouds reduce transmission at all wavelengths, leaving the ratios between wavelengths relatively unchanged. As shown in Methods based on data from [46], the ratio of light transmitted at wavelengths 400 nm and 660 nm changes by only 5% even with maximal cloud cover.

Humidity variations have a significant impact on spectrum as well [47, 48]. The distortions due to humidity primarily affect spectral ratios at the red of the spectrum while the blue end is relatively unaltered. As shown in Methods, based on the data of [48], a change in humidity from 20% to 80% can distort the ratio of red light to blue light by 50% but leaves the ratio of blue light to violet light virtually unchanged.

We see in Fig.3 that weather-related fluctuations can overwhelm the systematic variation in intensity over the lunar cycle but have a relatively smaller effect on spectrum. To produce these plots, we combined weather data[49] on cloud cover and humidity fluctuations for the Great Barrier Reef (July 2015) with data[45, 46, 48] on the impact of such fluctuations on natural light and measurements of the spectrum of natural light spectrum over the lunar cycle [44]; see Methods for more details.

**FIG 3.**
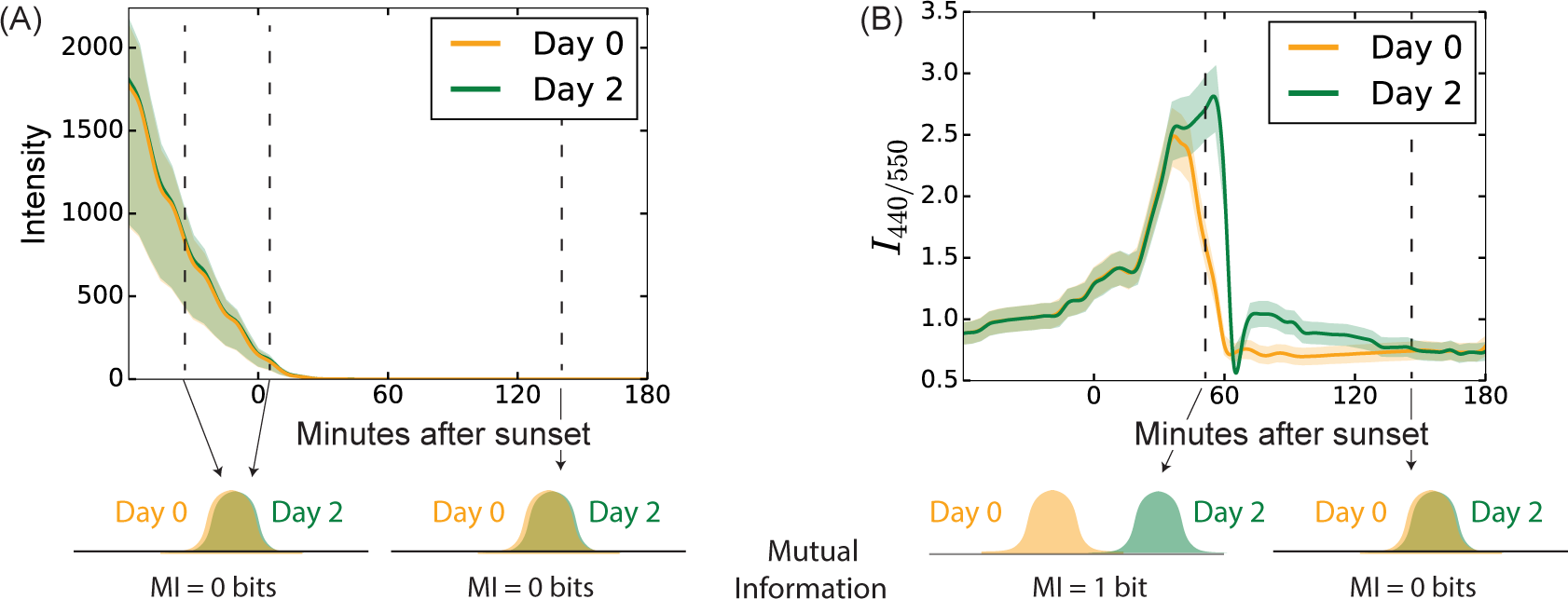
The spectrum of natural light is much more robust to atmospheric fluctuations than is intensity. (a) Cloud cover can change light intensity at the earth’s surface by orders of magnitude. As a result, Day 0 and Day 2 of the lunar cycle produce indistinguishable probability distributions of intensity when atmospheric fluctuations are included. (b) In contrast, the effect of cloud cover and humidity on spectrum is much smaller. Hence twilight spectrum on Day 0 and Day 2 are distinguishable at specific times (e.g., 55 minutes after sunset) despite atmospheric fluctuations. We quantify the distinguishability of probability distributions using Mutual Information. 1 bit of information reflects a perfect ability to distinguish a pair of days based on observed spectrum/intensity. (*I*_440*/*550_ in (b) is the relative photon catch in two opsins with peak sensitivity at 440 and 550 nm. Shaded regions in (a,b) are 1-sigma fluctuations, produced by combining weather statistics for the Great Barrier Reef in July 2015 [49] and data on cloud cover and humidity effects of natural light in [45, 46, 48])

### C. An information-theoretic approach to combine biophysical constraints and spectral and weather data

The circadian and circalunar regularities reported in Fig.2 are not likely to be biologically relevant if unpredictable atmospheric variations are larger than these systematic changes as shown in Fig.3. We therefore wanted to quantify the strength of biologically relevant signal given these natural perturbations in the spectral regularities of skylight, in a manner that does not make assumptions about specific biochemical processing that might occur inside an organism.

One approach is provided by Information theory [50, 51]. Information theory provides a rigorous way of determining the *highest* accuracy with which regularities can be perceived by an organism subject to a set of biophysical constraints.

We modified standard Information theory to account for data on downwelling twilight spectrum [17, 37, 44], spectral effects of humidity and cloud cover[46–48], weather fluctuations[49, 52] and biophysical constraints[7, 53] in the following way:

1. Twilight spectrum observations: We used recent experimental measurements [44] of the twilight spectrum in rural Pennsylvania with minimal artificial light pollution. By comparing data on different days of the lunar cycle, we developed a simple model that predicts the spectrum at any given time of twilight for any given lunar phase.
2. Monte Carlo simulation of weather: We obtained weather statistics [49, 52] quantifying the variation of cloud cover and humidity for the Great Barrier Reef (Australia) and for Death Valley (CA, USA). Using Monte-Carlo sampling, we simulated weather conditions with the same statistics as reported by these historical databases. We then applied the measured spectral and intensity distortions due to such weather (see Fig.3) to the signal predicted by the spectral model above. In this way, we populated a histogram of time courses of observed spectrum and intensity of natural light in these places over time under variable weather conditions.
3. Biophysical constraints: Organisms do not perceive time courses of natural light with perfect spectral and temporal accuracy. Instead, organisms might observe, e.g., the photon catch in a set of opsins, each with a specific spectral absorbance, integrated over an interval of time. Consequently, we projected the simulated high-dimensional histograms down to lower dimensional histograms that correspond to such biophysical constraints. We use these reduced histograms to compute the mutual information between, e.g., such spectral catches and the day of the lunar cycle.

The output of the Information theoretic calculation is in bits. In this case, 1 bit of information implies, for example, that the observed spectrum or intensity is sufficient to tell apart two specific days of the lunar cycle with complete confidence. See Fig.3 and Methods (Supplementary Fig. 2) for more information and a plot of this relationship.

In summary, our Information theoretic approach provides significant and complementary benefits: (a) it can combine data on twilight observations, spectral effects of clouds and humidity and historical weather data, (b) it can account for known biophysical constraints on receptor mechanisms, (c) it does not assume anything about how the signals are processed in the organism.

On the downside, Information theory can only provide an upper bound on the strength of biologically perceived regularities subject to constraints in (c); any particular organism can underperform relative to such a theoretical limit because of constraints not known and therefore unaccounted for.

### D. Impact of climate

The fluctuations shown in Fig.3 correspond to weather conditions at the Great Barrier Reef (July 2015) [49]. To see what kind of difference climate makes, we also ran our Monte-Carlo simulation based Information calculations on weather statistics corresponding to the Death Valley (CA, USA). The Death Valley has very different climate than the Great Barrier Reef - in particular, the fluctuations in cloud cover and humidity are significantly lower, especially in June.

Using these Monte-Carlo simulations, we computed the information available in intensity and spectrum to distinguish every pair of days *i, j* over the lunar cycle for the Death valley and the Great Barrier Reef. (Full moon is defined as Day 0.) See Fig.4. We find that the information in intensity is significantly enhanced at the Death Valley relative to the Great Barrier Reef due to the significantly lower cloud cover variations. Information in spectrum is also enhanced but to a much smaller degree. (As shown later, the information in spectrum can be significantly enhanced in all climates by having more than two opsins and integrating photon catch over specific intervals of the evening.)

**FIG 4.**
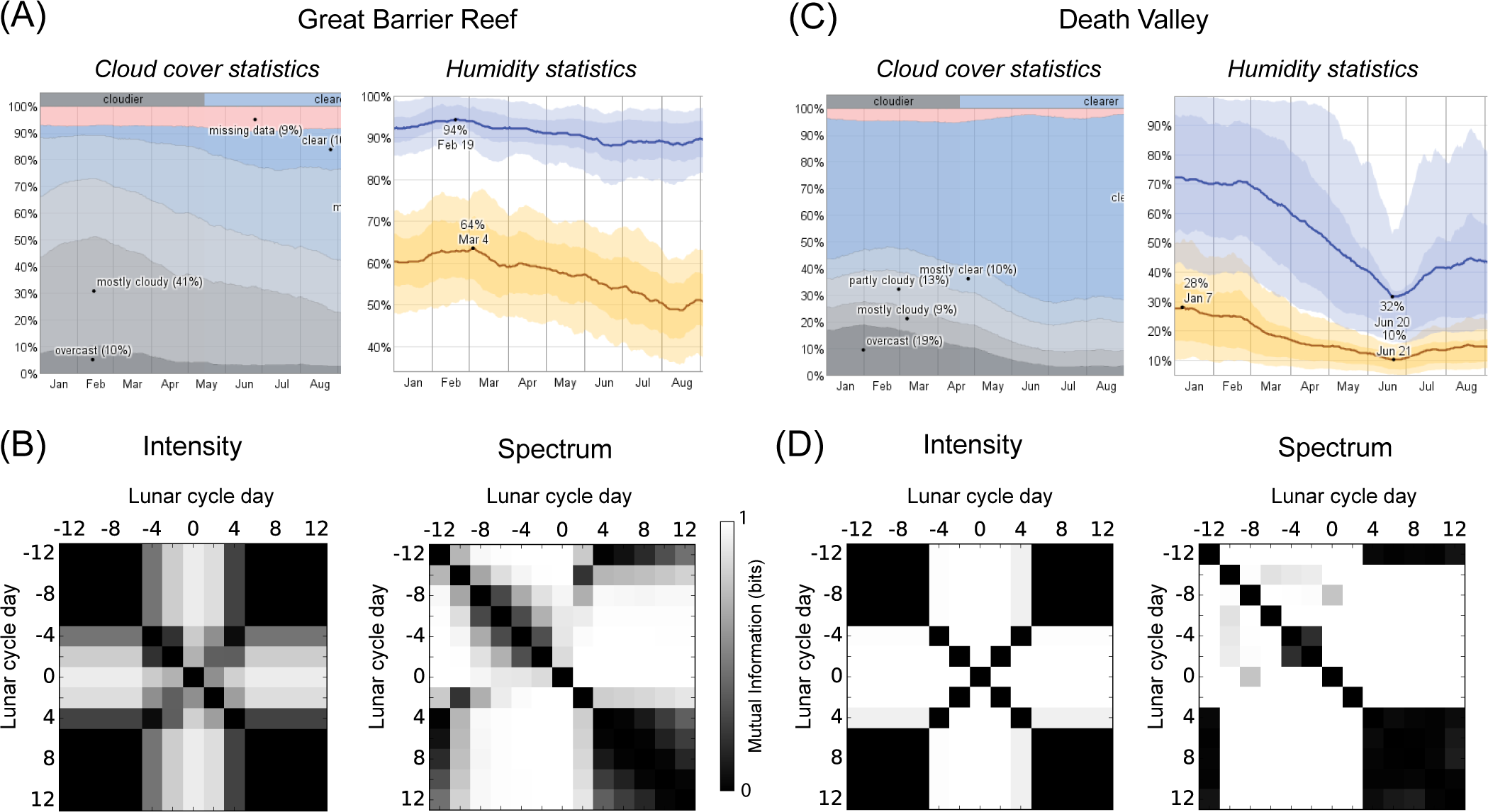
Climate has a large impact on information in intensity and a much smaller impact on information in spectrum. (a,c) Natural cloud cover and humidity variation are much higher near the Great Barrier Reef, Australia than in Death Valley, CA. We performed Monte-Carlo simulations of cloud cover and humidity variations using the weather statistics for these places [49, 52] and computed information from such simulations. (b) At the Great Barrier Reef, large cloud cover fluctuations impact moonlight intensity, making it impossible to distinguish most pairs of days over the lunar cycle. In contrast, the spectrum is robust to cloud cover and also humidity fluctuations. (Entry *i, j* of matrix shows available information in intensity (or spectrum) to distinguish day *i* and day *j* of the lunar cycle.) (d) With Death Valley-like weather, variations in cloud cover are much smaller and hence information in intensity is dramatically improved relative to the Great Barrier Reef. The spectrum shows smaller improvements. (Intensity plots reflect total light intensity over the night, spectrum plots reflect measurement at a specific time interval. See Methods for details.)

Thus, the information contained in spectrum is relatively robust to weather across a range of Earth’s climates, while the information contained in intensity is labile to weather and climate.

### E. Three opsins extract most available information

Terrestrial and marine species show a large range in the number of opsins with distinct peak sensitivities, possibly driven by gene duplication events followed by divergence[3, 10] The numbers range from three or four in several coral species [17] to six opsins in the sea urchin *Strongylocentrotus purpuatus* [54].

With multiple opsins, spectral information offers a distinct advantage that is not available in intensity - the information obtained by comparing multiple spectral “channels” (i.e., ratios) can be much greater than the sum of information carried by each of those channels. See Fig.5.

**FIG 5.**
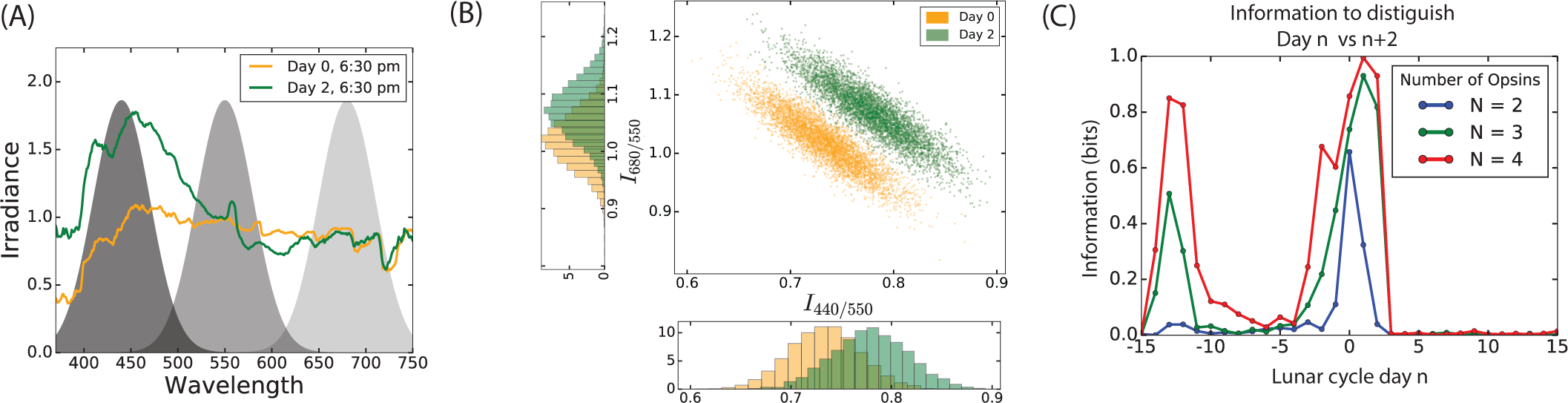
Signals from multiple opsins can be combined to increase information since atmospheric distortions have correlated effects across the spectrum. (a) Multiple opsins, each sensitive in a different wavelength range, allow an organism to measure multiple distinct spectral ratios. (b) Atmospheric distortions make it impossible to distinguish Day 0 (orange) from Day 2 (green) using any one of the two spectral ratios *I*_440_*/I*_550_ and *I*_680_*/I*_550_ (see overlapping 1-dimensional histograms). However, the distributions are easily separated in two dimensions, i.e., if both spectral ratios are measured at the same time. (c) Increasing the number of opsins from *N* = 2 to *N* = 3 significantly increases the available information about the lunar phase, while *N* = 4 opsins provide only a small additional benefit. Intuitively, one pair of blue and green-sensitive opsins allows one to ignore humidity fluctuations while a third opsin helps factor out cloud fluctuations. (Weather conditions and solar and lunar elevations corresponding to Great Barrier Reef, July 2015).

A particularly dramatic example of this phenomenon is presented in Fig.5b. The spectral signals on Day 0 and Day 2 are indistinguishable when viewed through *I*_680_*/I*_550_ or *I*_440_*/I*_550_ alone — see 1-dim histograms in Fig.5b. (Here, the spectral signal was collected between 45 and 110 minutes after sunset.) However, combining these two spectral channels — i.e., viewing the distributions in two dimensions as shown in Fig.5b — makes these distributions easily separable. Hence information in the sum of the spectral channels is much greater than the sum of the information in each individual channel in isolation.

Information theory allows us to expand such a comparison to more opsins and include time-series behavior. In Fig.5c, we systematically tested the information available in such combined spectral channels to discriminate between days across the lunar cycle for optimally positioned *N* = 2, 3, 4*, …* opsins. We find that going from *N* = 2 opsins to *N* = 3 provides a significant boost in information while *N* = 4 opsins provides a smaller benefit. Adding further opsins do not provide significant new information.

To understand this saturating effect around *N* = 3 opsins, note that if cloudy weather conditions change a given spectral ratio, say *I*_440_*/I*_550_, by some amount, the weather-related change in another ratio such as *I*_680_*/I*_550_ is entirely determined by this change (see Supplementary Fig. 1a). That is, the effect of cloud cover variations is not independent across the spectrum but rather constitutes only one independent source of variation.

Consequently, with only one independent source of spectral variation (e.g., cloud cover), two independent spectral ratios measured using *N* = 3 distinct opsins can potentially factor out cloud cover variations. Since our model has two independent sources of spectral variation (cloud cover and humidity), we expect that *N* = 4 opsins will provide significantly enhanced information. However, humidity variations do not affect spectral ratios in the blue end of the spectrum (see Supplementary Fig. 1b); hence *N* = 3 opsins already provides most of the available information if one pair of opsins is dedicated to the blue and green range of the spectrum.

Finally, we find that the exact positioning of *N* = 3 opsins does not significantly change information content, provided their sensitivities broadly cover the optical range as shown in Fig.5a, including one pair in the blue-green region that is relatively unaffected by humidity changes.

### F. Dynamic spectral signals and integrated photon catch by opsins

The rapid changes in spectrum at twilight could potentially be in conflict with the low intensity of light available to opsins in the brain or other such tissue. As shown in Fig. 3b, the spectral signal of downwelling sky-light, while robust and informative of lunar phase, is a highly dynamical signal that changes dramatically over the course of twilight[44]. The spectral signal is informative only in a specific interval of time. On the other hand, such opsins might need to integrate photon catch over many minutes to robustly resolve the spectrum.

To test the information available in integrated photon catches, we performed Monte Carlo simulations of the weather and collected the integrated photon catch with multiple opsins at different times of twilight and dawn-light. We find significant information even after integrating for up to 30 – 60 minutes. We find that the precise time interval capturing the ‘observation window’ strongly affects which days can be discriminated. As shown in Fig.6a, the interval 80 – 110 mins after sunset provides significant information towards discriminating days 0 – 2 of the lunar cycle while observations over the interval 200 – 230 mins are needed to distinguish days 2 – 5. Similarly, Fig.6b indicates that dawnlight observations over similar time intervals provide information about the days prior to the full moon (day 0). See also Supplementary Fig. 3 and 4.

**FIG 6.**
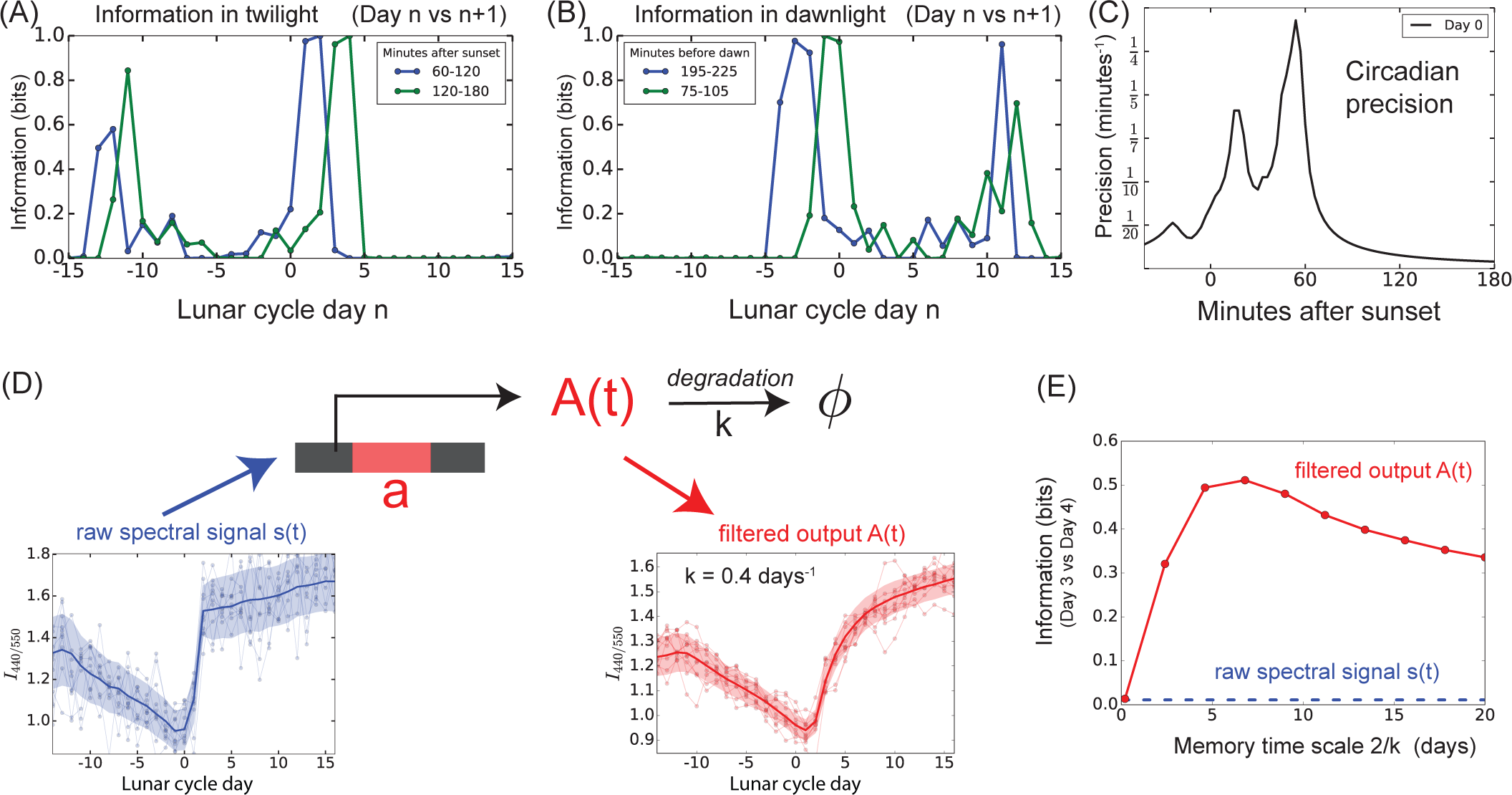
Integrating photon catch over different timescales offers different kinds of information because of the dynamic nature of spectral changes. (a,b) Integrating photon catch over different intervals of twilight and dawn hours carries information about different parts of the lunar cycle. (c) The rapid changes in spectrum at twilight also provides a sharp estimate of the time of the day relative to sunset. Such circadian time-telling precision is particularly high (±4 mins) about 60 minutes after sunset. (d) Multi-day memory can greatly enhance lunar phase information. Expression of protein *A* (red) is promoted by the spectral signal *s*(*t*) = *I*_440_*/I*_550_ (blue); *A* then degrades on a slow multiday timescale ∼2*/k*. (e) Protein level *A*(*t*) (red) is less noisy and hence significantly more informative about lunar phase than the raw spectral signal *s*(*t*). Information is maximized at a specific degradation timescale 2*/k* 5 days; shorter memory is unable to filter weather fluctuations while longer memory averages out the systematic spectral changes over the lunar cycle. (Simulated weather conditions corresponding to the Great Barrier Reef, July 2015 [49].)

### G. Twilight spectral changes carry information about time of day

Within a single 24-hour circadian cycle, the spectral signal shows a sharp feature at dawn and at evening twilight that can be used as a time-telling device. Traditionally, the information-bearing signal for circadian time is assumed to be intensity, since intensity of natural light does change dramatically over twilight hours. However, the precision and reliability of such time telling is lower when weather fluctuations are taken into account.

To quantify the precision of circadian time-telling based on spectral signals, we computed the systematic changes in the mean level of the spectral signal *I*_440_*/I*_550_ over the course of twilight hours as well as the variations due to simulated atmospheric fluctuations (corresponding to the Great Barrier Reef). We define precision of time-telling to be the (reciprocal of) one standard deviation error *δt* in the time deduced from the value *I*_440_*/I*_550_ due to atmospheric fluctuations. We find that the error *δt* is as low as ∼4 minutes around 60 minutes after sunset on day 0 of the lunar cycle (the night of the full moon). See Fig.6c. Thus spectrum of twilight can provide a highly reliable, stereotyped signal at a specific circadian time. Such a signal has been proposed as a cue for synchronized reproductive behaviors in marine organisms such as corals [17].

### H. Multi-day memory can filter weather fluctuations and increase lunar phase information

Thus far, we have quantified the ability of an organism to infer the lunar phase based on natural light observed over the course of one evening (or dawn or night), assuming no memory of light received on prior days.

However, the internal biochemistry of an organism can likely retain some memory of the signal over multiple days. Such a memory of the signal seen over many days offers more information than a snapshot seen on a given day since weather changes can potentially be averaged out.

We show that information is indeed enhanced by memory of a specific timescale using a simple biological realization of such cellular memory. In Fig.6d we consider expression of a gene product or membrane potential stimulated by a specific spectral signature, followed by a slow degradation of that product or potential on a timescale 1*/κ* that varies from under a day to multiple days. We then compute the mutual information between protein level *A*(*t*) on a given day and the lunar phase of that day. This information captures how well the concentration or magnitude of *A* can be used to determine the lunar phase.

We see that the time courses of the output *A*(*t*) in Fig.6d, is significantly less noisy than the input *s*(*t*). Consequently, we find that the information in *A*(*t*) to distinguish Day 3 and Day 4 of the lunar cycle can be significant, although these days are nearly indistinguishable based on the raw input signal *s*(*t*).

To understand such information enhancement intuitively, note that *A*(*t*) effectively reflects the moving average of the natural light signal *s*(*t*) over a timescale ∼1*/k*. Averaging for too short a period (i.e., large *k*) is not effective at averaging out weather-related fluctuations. However, averaging for too many days also results in low information, since the systematic change in the spectrum over the lunar cycle is also washed out along with weather-related fluctuations. In fact, we find that the information in *A*(*t*) is highest with a memory timescale of 2*/k* ∼ 5 days (see Fig.6e).

Biology can, in principle, build more complex machinery with a more complex memory structure that e.g., retains memory of the signal at a specific time in the past. However, our results suggest that even the simplest forms of memory of the right timescale can strongly enhance regularities in the relative photon catches of multiple opsins.

## III. DISCUSSION

Non-imaging opsins have been found in an increasing number of tissues in recent years [8, 9]. Their biological role is being actively sought [8, 11–15] and early evidence suggests that disruption does have functional consequence. While intensity of light is obviously lower inside tissue than at the surface of an animal’s body, even a decrease of 6 log units would still leave sufficient flux within the tissue to be physiologically relevant, especially if the cells involved potentially have long integration times and long memories[22, 55], as we predict here.

Our work focuses on the stable reliable patterns in the spectrum of such natural light - the ‘natural scene’ for such non-imaging systems - that has been present over evolutionary timescales. Stable patterns in ‘natural scenes’ are known to be exploited in similar sensory contexts, ranging from edge detectors in imaging vision[29– 32] to blue-sensitive opsins behind the retina[3, 5, 13] that exploit natural light rhythms to set circadian rhythms. In addition to coupling to the regular periodic patterns in natural light quantified here, non-imaging opsins are also likely to have roles in sensing intermittent light from the local environment [3].

While ideas about the functional role of non-imaging opsins are currently speculative, many reproductive behaviors synchronized to the lunar cycle are already well-documented. For example, the mass spawning in corals and other marine organisms is known to be linked to the lunar phase, with each species releasing gametes en masse in a distinct narrow window of the lunar cycle. Such synchronized release of gametes is thought to increase the chances of fertilization. Numerous studies have sought to identify mechanisms that trigger such a lunar cycle synchronized response[28]. Since these organisms do possess multiple opsins, natural light is suspected to serve as a cue [17, 55]. Spectral signals at twilight are also thought to affect foraging behavior of various terrestrial species [56, 57].

While circalunar behaviors are seen in numerous organisms, free running circalunar clocks have been found in only a few organisms [41]. However, our results suggest that a complex circalunar clock is unnecessary in organisms with multiple opsins. A simple response strategy to the spectral signal, possibly combined with a simple filtering mechanism like that described in Fig.6d, is sufficient to show circalunar behavior with timing precise to a single day within the lunar cycle.

Our work also suggests spectral changes as an alternative *zeitgeber* for circadian behavior, one that is significantly more reliable than light intensity alone, which is usually taken as the driver of circadian behaviors [58]. Recent work suggests that intensity fluctuations in light do affect the quality of circadian machinery in photosynthetic organisms[59]. Organisms with multiple opsins could avoid such costs by accessing the more reliable spectral signal. Indeed, such spectral changes have been proposed as a cue for some precise circadian behaviors seen in marine organisms [17, 22, 37] that coincide with twilight hours.

In summary, our work on information on the ‘natural scene’ for non-visual opsins fills in a gap in natural signals available at a timescale between daily and yearly cycles. While numerous entraining signals are known for daily and yearly biological rhythms, few such universally available reliable signals are known for the ∼28 day lunar cycle. Much as the amount of light (e.g., photoperiod) changes over the year and is implicated in many important biological rhythms such as flowering [60], our work suggests that the spectrum of natural light provides reliable information about the lunar cycle. These mechanisms could also be relevant to reproductive timing in mammals including humans, where emerging data suggests that there are fitness costs associated with off-season births[43, 61, 62]. Our information theoretic approach quantifies the reliability of such a signal with minimal assumptions and yet is readily tailored to specific climates and biophysical constraints in specific organisms.

## IV. METHODS AND MATERIALS

### A. Spectrum

We use the data of [44] to build a model of the the downwelling spectrum *S*(*λ*) as a function of lunar phase and the time of the day.

We begin by modeling the spectrum as a linear combination of the contributions of the sun and moon, where the contribution of the moon is modulated by its phase *ϕ*:

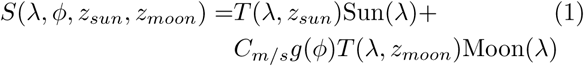

with all quantities defined in the table I.

**TABLE 1.**
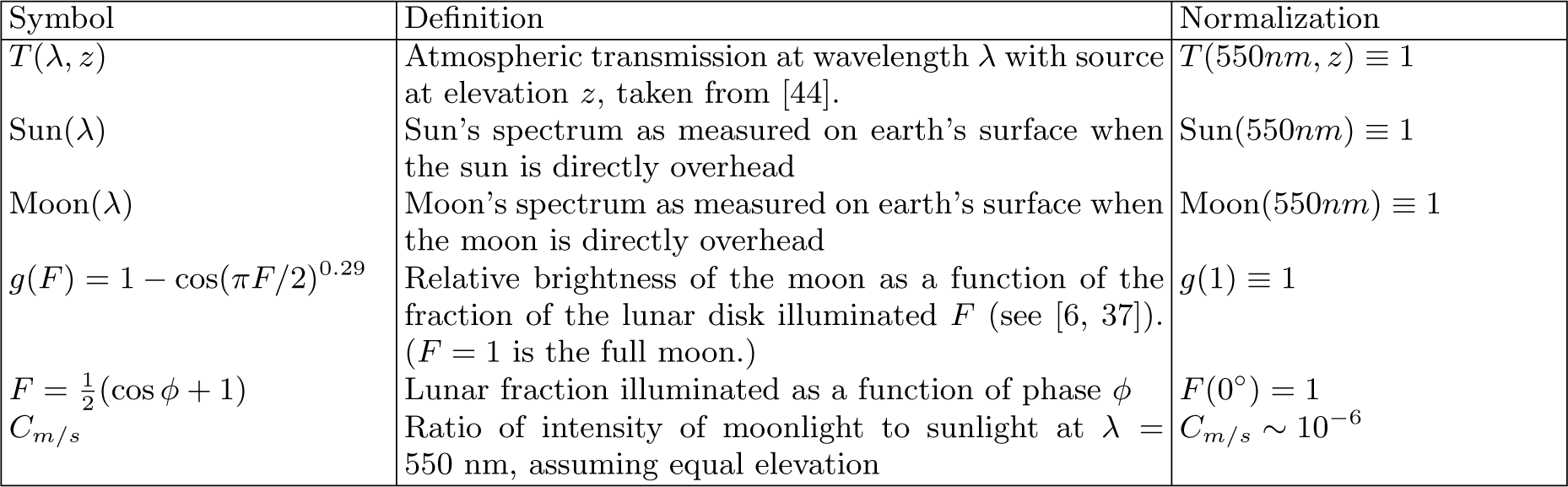
Quantities used in the spectral model that was fit to experimental measurements of twilight spectral in [44].

| Symbol | Definition | Normalization |
| --- | --- | --- |
| $T(\lambda, z)$ | Atmospheric transmission at wavelength $\lambda$ with source at elevation $z$, taken from [44]. | $T(550nm, z) \equiv 1$ |
| $\text{Sun}(\lambda)$ | Sun's spectrum as measured on earth's surface when the sun is directly overhead | $\text{Sun}(550nm) \equiv 1$ |
| $\text{Moon}(\lambda)$ | Moon's spectrum as measured on earth's surface when the moon is directly overhead | $\text{Moon}(550nm) \equiv 1$ |
| $g(F) = 1 - \cos(\pi F/2)^{0.29}$ | Relative brightness of the moon as a function of the fraction of the lunar disk illuminated $F$ (see [6, 37]). ($F = 1$ is the full moon.) | $g(1) \equiv 1$ |
| $F = \frac{1}{2}(\cos \phi + 1)$ | Lunar fraction illuminated as a function of phase $\phi$ | $F(0^\circ) = 1$ |
| $C_{m/s}$ | Ratio of intensity of moonlight to sunlight at $\lambda = 550$ nm, assuming equal elevation | $C_{m/s} \sim 10^{-6}$ |

Refs [44] recently measured the spectrum of down-welling radiation over the course of the day and night, including twilight hours, on different days of the lunar cycle.

We used non-linear regression to fit our model above to the data in [37, 44] to determine Sun(*λ*), Moon(*λ*), *C*_*moon/sun*_.

This model then serves as the signal to which we add variations due to weather and atmospheric conditions below. To find the signal on a day of lunar phase at a given time t of the day, we calculate

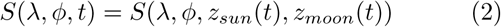

where *z*_*sun*_(*t*)*, z*_*moon*_(*t*) are the elevations of the sun and moon at time t. We obtain these elevations along with the corresponding lunar phases for various locations and times from charts available at [63].

### B. Atmospheric distortions

#### 1. Effect of clouds

We use a model of the spectral effects of clouds from [46], which gives the cloud-distorted spectrum *S*_*cloud*_(*λ, t*) = *C*(*λ, f*)*S*(*λ, t*), where *S*(*λ, t*) is the spectrum with clear skies taken from Eqn.2 and the spectral cloud effect

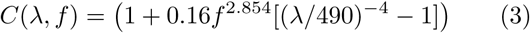

gives the factor by which irradiance is reduced at wavelength *λ* and cloud fraction *f* (see also Supplementary Fig. 1a).

#### 2. Effect of humidity

We use data from [48] to model the effect of relative humidity according to

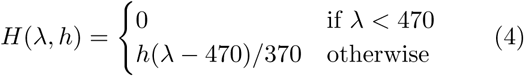

where *h* is the relative humidity and *H* gives the humidity-distorted spectrum by *S*_*hum*_(*λ, h*) = *H*(*λ, h*)*S*(*λ*), with *S* the spectrum at 0% relative humidity, taken from Eqn.2 above (see also Supplementary Fig. 1b).

#### 3. Intensity variations with cloud cover

We model the effect of clouds on the intensity of down-welling light according to [45] and [46]. We calculate the mean fractional reduction in intensity for cloud fraction f from [46] according to *C*(*f*) = 1 *−* 0.674*f* ^2.674^, where *C*(*f*) = *S*(*f*)*/S*(*f* = 0) is the ratio of light intensity for cloud fraction *f* to clear-sky intensity *f* = 0.

### C. Weather Statistics by Location

We use weather data from weatherspark.com to model the distribution of fractional cloud cover (fraction of sky obscured by clouds) and relative humidity at the Great Barrier Reef and in Death Valley, CA for the months of June and July. For each location and form of variation we approximate the distribution as Gaussian, truncated at 0 and 1, with mean and sigma obtained from the on-line weather data. For the reef, we use mean *µ*_*c*_ = .65, standard deviation *σ*_*c*_ = .3 and mean *µ*_*h*_ = .6, *σ*_*h*_ = .2 for cloud cover and percent humidity respectively [49]. For Death Valley, we use *µ*_*c*_ = 0, *σ*_*c*_ = .2 and *µ*_*h*_ = .25, *σ*_*h*_ = .2 [52].

In Fig.3a we plot total intensity under average atmospheric conditions at the Great Barrier Reef, integrated over three minute intervals for Day 0 and Day 2 (with Day 0 being the day of the full moon). In Figure 3b we plot the ratio of light absorbed by opsins with peak sensitivity at 440 nm to light absorbed by opsins peaked at 550 nm under average Great Barrier Reef weather conditions, again integrated in three minute intervals for Day 0 and 2. In each plot, the shading shows the variation in the signal at plus and minus one standard deviation in atmospheric conditions. Solar and lunar elevation data are from the Great Barrier Reef in July 2015. In Fig.4 we sample 10,000 sets of atmospheric conditions typical of the Great Barrier Reef (a,b) or Death Valley (c,d) and calculate the total light intensity (red) and opsin catch ratio *I*_440_*/I*_550_ (blue) for each set for a given measurement strategy. In Fig. 4b we plot the pairwise mutual information between days (i,j) (*i* ∈ (1, 2*, …,* 28)) and intensity and between days and spectrum. The intensity signal for each day was obtained by integrating from 8:30 pm - 1:30 am (*-*35° to *-*70° solar elevation) with start and stop times drawn from a Gaussian distribution with a standard deviation of 5 minutes. The spectral signal was obtained by the ratio of opsin catches *I*_440_*/I*_550_ and *I*_680_*/I*_550_, each integrated from 6:30 pm to 7:30 pm (*-*10° to *-*25° solar elevation), with start and stop times drawn from a Gaussian distribution with a standard deviation of 5 minutes. All solar and lunar elevations are drawn from data at the Great Barrier Reef in July 2015. Figure 4d follows the same pattern but with solar and lunar elevations and typical atmospheric conditions from Death Valley in July 2015. Measurement times are adjusted so that solar elevations are consistent across the Death Valley and Great Barrier Reef plots.

#### Other sources of spectral variation

Diffuse downwelling irradiance can be split into three components [47]: Rayleigh scattering, Mie scattering and a component due to multiple reflection of light between the ground and atmosphere, which varies with albedo [47]. Rayleigh scattering, or scattering due to particles with radii much smaller than the wavelength of scattered light, is responsible for the changes in the color of light with solar elevation. Mie or aerosol scattering is characterized by scattering from particles whose radii are the same order of magnitude as the wavelength of scattered light.

For the purpose of this paper, we are primarily concerned with the temporal variation of the spectrum at some fixed geographical location. Our regression-based model of the spectrum is generated from data collected in rural Pennsylvania [44]. In other regions, factors such as ozone transmittance, the presence of dust and surface albedo may produce different mean values of irradiance. However, we assume that the first order *variations* in downwelling irradiance can be captured across a range of locations by modeling the reduction in irradiance as a function of fractional cloud cover and percent humidity.

### D. Biophysical constraints

#### 1. Opsin details

We modeled the spectral sensitivity of each opsin type as a Gaussian with variance of 90 nm and a mean of 440 nm, 550 nm or 680 nm. These values were chosen to roughly maximize information retrieval across the visual spectrum for a range of integration periods. The exact optimal positioning depends on the times or days to be distinguished. However, we found the ability to measure variations in irradiance across the visible spectrum to be more important for extracting available information than the precise placement of the peak absorption wavelengths. Note that in reality, opsin absorption spectra tend to have a longer tail at higher wavelengths, with the overall variance increasing with the maximum absorption value [53].

In Fig. 5a, we plot irradiance integrated over 1 hour (6:30 pm - 7:30 pm, or −10° to −25° solar elevation) in 1 nm bins, assuming clear skies and 0 percent relative humidity. Solar and lunar elevation data are from the Great Barrier Reef in July 2015. The shaded regions are sample opsin absorption curves peaked at 440, 550 and 680 nm. In Fig. 5b we plot 5,000 opsin catch ratios, with weather conditions drawn randomly from our calculated distribution for the Great Barrier Reef. The ratios are obtained by integration from 6:30 - 7:30 pm (equivalently, −10 ° to −25° solar elevation). Solar and lunar elevation data are from the Great Barrier Reef in July 2015. In Fig. 5c we plot the information content of days (n, n+2) for n = −15, −14… 0,..,15, or from one new moon to the next. The lines correspond to information obtained by 2, 3, or 4 opsins with peaks at (440 nm, 550 nm), (440 nm, 550 nm, 680 nm) and (440 nm, 550 nm, 620 nm and 680 nm) respectively. In each case, irradiance is integrated from approximately 6:30 pm - 7:30 pm (−10° to -25° solar elevation), with start and stop times drawn from Gaussian distributions with a standard deviation of 10 minutes. Solar and lunar elevation data are from the Great Barrier Reef in July 2015.

#### 2. Integrated photon catch

We consider a raw spectral signal *S*(*t, λ*), which is then distorted according to atmospheric conditions. The distorted signal *S*_*d*_(*t, λ*) is convolved with the opsin absorption curves *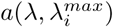* and integrated over a period of time to give the photon catch

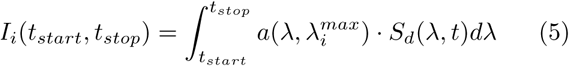

. Here *i* = 1, 2*…N* are the *N* opsins with peak absorption at 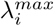, where 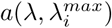 is a gaussian with *σ* = 90 nm centered at 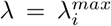. We then choose one opsin channel to normalize against (take it to be the *N* th opsin) and calculate the measured catch ratios *R*_*i*_ = *I*_*i*_*/I*_*N*_ for *i* = 1, 2..*N-*1.

In Figure 6a and b, we plot the information content of days (n, n+1) for n = −15,…0,…15, with information obtained by the integration of opsin catches at 440 nm and 550 nm for two 30 minute periods each in evening (a) and morning (b). Start and stop times are drawn from a Gaussian distribution with a standard deviation of 5 minutes.

#### 3. Circadian precision

For a given set of opsins and visual strategy, we define the circadian precision as 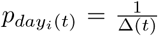, where Δ(t) is defined as the smallest time Δ such that the distribution of catch ratios *R*_1_ = *R*(*t*) and *R*_2_ = *R*(*t* ±Δ) are more than one standard deviation removed from each other; that is, assuming without loss of generality that *R*_1_ *< R*_2_, Δ(*t*) is the smallest time for which *R*_1_ + *σ*(*R*_1_) *< R*_2_*-σ*(*R*_2_).

In Fig.6c, we plot the circadian precision for Day 0 and Day 2 assuming opsins peaked at 440 nm and 550 nm, and atmospheric conditions characteristic of the Great Barrier Reef. We bin time in three minute intervals, such that each measured ratio *R*_*i*_(*t*) = *I*_*i*_(*t*_*start*_*, t*_*stop*_)*/I*_*N*_ (*t*_*start*_*, t*_*stop*_) is obtained according to (5) with *t*_*start*_ = *t-*3 minutes and *t*_*stop*_ = *t* minutes. Solar and lunar elevation data are from the Great Barrier Reef in July 2015.

#### 4. Biochemical temporal filtering

We consider the effect of simple biochemical mechanisms with multi-day memory of photon catches. Such mechanisms can act as low pass filters that reduce the impact of atmospheric fluctuations, provided the filtering timescale is tuned appropriately. We performed these simulations in the following way: to obtain the signal for Day n, we repeatedly sample trajectories, representing random weather patterns, over many days before Day Each of these generates a noisy signal *S*(*λ, t*), from which we calculate the opsin catch ratios *R*_*i*_ = *I*_*i*_*/I*_*N*_, with *I*_*i*_*, I*_*N*_ as defined in (5).

One ratio *R* is then fed into the circuit shown. The circuit is defined by the following equation:

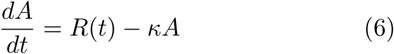

where *R*(*t*) is updated every 24 hours. Thus, the signal *R*(*t*) promotes the expression of *A* which degrades on a time scale *κ*. This implies

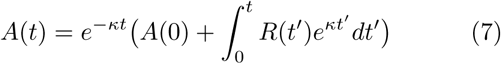

For each weather trajectory ending in Day n, we obtain one value of the signal on Day n, defined as the concentration of *A* on Day n, or *A*(*t* = *n*).

In Fig. 6d-1, we plot a sample of ratios *R*(*t*) obtained by integrating opsin catches at 440 nm and 550 nm from approximately 6:30 pm - 7:30 pm under varying atmospheric conditions. Start and stop times are drawn from a Gaussian distribution with a standard deviation of 5 minutes. In Fig. 6d-2 we plot the moving averages *A*(*t*) of the sampled signals as defined in (7), with *κ* = 0.4. Figure 6e shows the information content of Days 3 and 4 when passed through the filter for varying values of *κ*, with the same visual strategy as in 6 (d). Solar and lunar elevation data are from the Great Barrier Reef in July 2015.

### E. Computing Mutual Information

#### 1. General formula

The main quantity we compute repeatedly in this paper is the mutual information between lunar phase (equivalently day of the month) and observed spectral data. Given a pair of days with phases *ϕ*_*a*_ and *ϕ*_*b*_, we compute

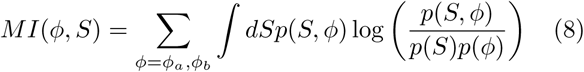

where *S*(*λ, t*) is the set of all possible spectra for the corresponding day *a* or *b* under varying atmospheric conditions, and *p*(*S, ϕ*) represents the joint probability of observing the spectral signal *S* while the moon is in the phase *ϕ*.

We write the joint distribution *p*(*S, ϕ*) = *p*(*S|ϕ*)*p*(*ϕ*) and set a uniform prior *p*(*ϕ*_*a*_) = *p*(*ϕ*_*b*_) = .5 to obtain

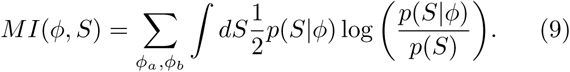

The value given by (9) represents the amount of information about the phase contained in (one’s measurement of) the spectrum, or the degree of certainty with which one can tell the phase when given only the spectrum.

If the observed spectrum is always sufficient to distinguish the two lunar phases with complete confidence, the mutual information between the spectrum and lunar phase will equal one. If measuring the spectrum is no use in determining the day – if someone attempting to infer the day from the spectrum does no better than guessing – then their mutual information is zero.

#### 2. Monte-Carlo sampling to mimic weather and temporal biophysical constraints

We determine the probability distributions 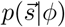 by Monte-Carlo sampling of a range of spectra, with mean determined by the regression model and variance determined by modeled fluctuations in cloud cover and humidity.

For a given location, we take a random sample of cloud cover fractions from the distribution determined by weather data, which is assumed constant over several hours. For each cloud cover *f*, we calculate the distorted spectrum *S*_*d*_(*λ, t*) = *C*(*λ, f*)*H*(*λ, f*)*S*_*clear*_(*λ, t*), where *C*(*λ*) and *H*(*λ*) are the fractional reduction in clear sky irradiance at cloud fraction *f* and relative humidity *h*, as defined in 3, 4 respectively. The values of *f* and *h* are held constant over the period (*t, t* + *δt*) for which the signal is integrated. In other words, they represent average weather conditions on a given day or night.

We generate *S*_*d*_ for a range of atmospheric conditions and then bin by irradiance, with each histogram axis corresponding to a distinct time and absorption peak 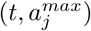. We calculate the probability of measuring a photon catch in the ranges *s*_*j*_ for times (*t*_1_*…t*_*n*_) ∈ *T* and opsin sensitivities (*a*_1_*…a*_*m*_) ∈ *A* as

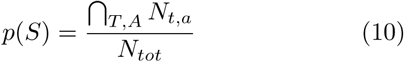

where *N* is the number of values in the *s*_*j*_ bin on each axis (*t, a*), and *N*_*tot*_ is the total number of simulated spectra for the given day. We set a fixed bin width across all histograms. To ensure that the histogram accurately reflects the distribution of spectra, the number of simulated spectra is chosen to be the square of the total number of bins. For calculations involving intensity, we set a minimum bin size corresponding to the resolution limit of non-imaging eyes, based on data from [7].

We include a small uncertainty in the timing of temporal measurements. For a given integration period (*t, t* + *δt*), we take the start and stop times of integration to be normally distributed, with a standard deviation *σ*_*t*_ ≈ *t/*6.

#### 3. Gaussian approximation

For calculations with many snapshots and many opsins, the computation above becomes intractable. For example, with 4 snapshots and 3 opsins, one has 12 channels. Populating a 12-dimensional histogram and then computing mutual information from that is not practical. Hence, for high dimensional cases we make a Gaussian approximation to this histogram. If the signal is *s*_*i*_, we take *N* = 10^4^ weather samples and create a data matrix *X*_*ai*_ of size *N* ×length 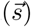. We then find the Gaussian that best approximates this distribution by computing the mean *µ* _*i*_ = ∑ _*a*_*X*_*ai*_ and variance 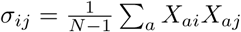.

We estimate the MI for this Gaussian model as follows. We will use the relation

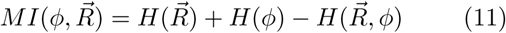

For a given pair of days *a* and *b*, we treat each as equally like to occur, so

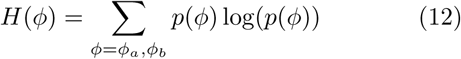

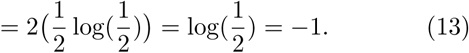

To calculate 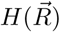, we draw another *N* = 10^4^ random samples from the Gaussian distributions for days *a* and *b*, and calculate the resulting catch ratios *R*_*i*_ = *I*_*i*_*/I*_*N*_ according to 5. We then write

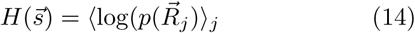

where the average is over all catch ratios 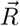 calculated from the spectral distributions 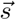 of Day 1 and Day 2. The probability 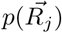 is obtained by 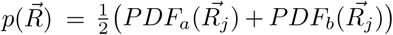 where PDF_*a,b*_ are the probability distribution functions of the Gaussians for days *a* and *b*.

To calculate the joint entropy, we write

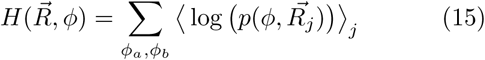

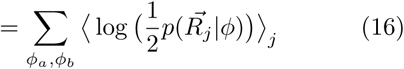

where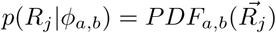

The mutual information is then calculated according to equation 11.

## ACKNOWLEDGEMENTS

We thank Sofia Magkiriadou, Stephanie Palmer and Michael Rust for insightful discussions. AM is grateful to the Simons Foundation MMLS investigator program for support. We acknowledge the University of Chicago Research Computing Center for computing resources.

